# Polymer physics models reveal structural folding features of single-molecule gene chromatin conformations

**DOI:** 10.1101/2024.07.16.603769

**Authors:** Mattia Conte, Alex Abraham, Andrea Esposito, Liyan Yang, Johan H. Gibcus, Krishna M. Parsi, Francesca Vercellone, Andrea Fontana, Florinda Di Pierno, Job Dekker, Mario Nicodemi

## Abstract

Here, we employ polymer physics models of chromatin to investigate the 3D folding of a 2Mb wide genomic region encompassing the human *LTN1* gene, a crucial DNA locus involved in key cellular functions. Through extensive Molecular Dynamics simulations, we reconstruct in-silico the ensemble of single-molecule *LTN1* 3D structures, which we benchmark against recent in-situ Hi-C 2.0 data. The model-derived single molecules are then used to predict structural folding features at the single-cell level, providing testable predictions for super-resolution microscopy experiments.

## INTRODUCTION

Recent DNA technologies, such as Hi-C [1,2], GAM [3,4] and SPRITE [5,6], have shown that mammalian chromosomes have complex, non-random 3D architectures within the cell nucleus, encompassing multiple folding structures across genomic scales [7–11]. Such an organization includes, for example, DNA loops [12], Topologically Associating Domains (TADs) [13,14], metaTADs [15], and larger architectural features, such as A/B compartments [1], lamina-associated interactions [16] and nuclear chromosome territories [17]. These chromatin structures play critical roles in gene regulation, as distal DNA regulatory elements (e.g., enhancers) have been reported to establish specific physical contacts with their target genes, particularly within TADs, to orchestrate cell transcriptional programs [18–23]. Misfolding of chromatin 3D structure, which can lead to abnormal gene-enhancer interactions, has been indeed implicated in genetic disease [24–27].

Moreover, advancements in super-resolution microscopy techniques have enabled the direct visualization of chromatin conformations at nanometer scales within individual cell nuclei, providing quantitative insights on single-cell genome structures [28]. These methods have revealed, for instance, that TAD-like domains exhibit spatially segregated globular conformations in single cells, albeit with significant cell-to-cell variability, highlighting crucial constraints on chromatin folding at the level of single DNA molecules [29–36].

On the other hand, theoretical models of chromatin have been instrumental in bolstering experimental technologies to understand the 3D organization of the genome [37]. These models, including polymer-physics-based and computational approaches [38–80], have been used, e.g., to dissect the fundamental molecular mechanisms shaping DNA contact formation and to derive quantitative testable predictions particularly in disease contexts [81–84].

In this work, we focus on a reference polymer physics approach, the Strings and Binders (SBS) model of chromatin [68,69], which has been extensively validated against independent experimental datasets, such as bulk Hi-C, GAM, and single-cell microscopy data [85–89]. In the SBS picture, physical contacts between distal DNA sites (such genes and enhancers) are established by diffusing cognate binding factors, which can bridge DNA binding sites through thermodynamic mechanisms of phase transitions [40,87]. Here, we apply the SBS model to investigate the single-molecule folding of a 2Mb wide genomic locus (Chr21:28-30Mb, hg38) encompassing the *LTN1* gene in human WTC-11 cells (a human induced pluripotent stem cell line), a crucial locus involved in diverse cellular functions ranging, e.g., from embryonic development and targeting of misfolded proteins to the onset of neurodegenerative diseases [90–92].

By performing massive Molecular Dynamics (MD) simulations of the model, we derive an ensemble of in-silico 3D conformation of the gene locus that we validate against recent in-situ Hi-C 2.0 data, generated via a highly optimized Hi-C protocol [93], available from the 4DN Data Portal (dataset reference number 4DNESJ7S5NDJ) [94,95]. Next, by leveraging the model single-polymer conformations, we conduct structural analyses at the single-molecule level, including spatial distance matrices, assessment of cell-to-cell folding variability, and 3D shape descriptors, producing non-trivial predictions that can be tested by real single-cell microscopy (e.g., super-resolution multiplexed FISH) experiments.

Overall, the model provides a validated, quantitative blueprint for assessing the spatial organization of key human genomic loci at the single-molecule level, complementing empirical investigations in the understanding of chromatin structures in single cells.

## METHODS

### The Strings and Binders (SBS) polymer model

The Strings and Binders (SBS) polymer model envisages a theoretical framework for understanding the 3D organization of chromatin where molecular interactions between distant genomic regions are driven by diffusing agents, such as Transcription Factors (TFs) or coactivators, that diffuse in the nuclear environment [68,69]. In the model, a chromosomal segment is represented as a coarse-grained, self-avoiding walk (SAW) polymer chain with specific binding sites for molecular bridging binders (a schematic model cartoon is shown in **Figure 1a**).

**Figure 1.**
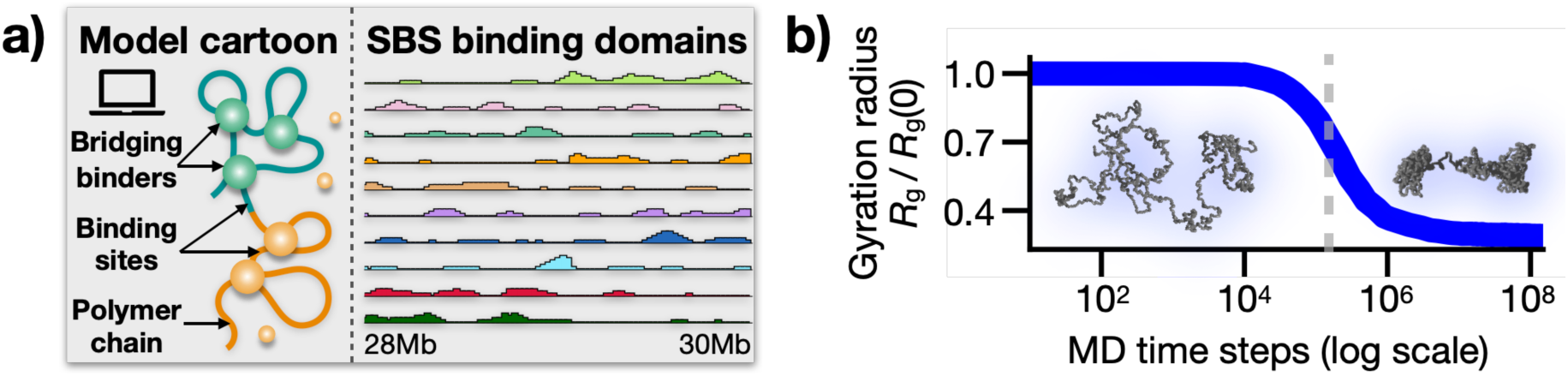
**a)** In the SBS model, a chromatin region is represented a self-avoiding polymer chain along which specific binding sites are arranged for diffusing, cognate molecular binders. By bridging cognate sites on the chain, the binders drive the folding of the polymer forming microphase-separated globular structures. The SBS binding domains of the studies *LTN1* locus (Chr21:28-30Mb) in human WTC-11 cells are shown along with a schematic cartoon of the polymer model. **b)** The polymer gyration radius *R*_g_ is shown here as a function of the MD time iteration steps (y-axis normalized by the *R*_g_ value at *t*=0). The function exhibits a sharp drop around 10^5^ time steps, signaling the collapse of the chain from an initial coil (i.e., randomly folded) to an equilibrium globule conformation [96]. Representative coil and phase-separated globule 3D structures are shown, respectively, below and above the phase transition point (grey shaded line in the figure).

In the simplest case of an SBS homopolymer chain, where the binding sites are all equal, the model has a phase transition from an extended coil (randomly folded) to a compact globular state [96] as soon as the binder molar concentration or their attractive interaction strengths exceed critical thresholds. For weak biochemical affinities (i.e., units of K_B_T), and for genomic resolutions approaching the sub-megabase scale, these thresholds are in the range of a few micromole/l [40,87], consistent with typical TF concentrations observed in-vitro [97].

To dissect the folding dynamics of real chromatin regions characterized by complex genomic contacts, the SBS model can be extended to include multiple types of binding sites, each associated to a specific type of cognate molecular binder [40]. Such heteropolymer configurations induce micro-phase separations of the polymer chain, leading to the formation of distinct globular domains enriched with specific binding motifs [87]. The features of these binding sites (i.e., their genomic location and relative abundance) are determined through a previously published machine learning approach, named PRISMR [83], which employs standard Monte Carlo-based optimization procedures to infer the optimal (i.e., minimal) SBS model of specific chromatin regions based solely on input bulk (e.g., Hi-C) contact data, with no additional fitting parameters. In our studied *LTN1* locus in human WTC-11 cells (Chr21: 28-30Mb), PRISMR returns a polymer of 800 beads with ten distinct types of binding sites, visually represented by different colors (**Figure 1a**). Interestingly, these domains have an overlapping, combinatorial organization along the chain, which has been shown to be crucial to explain chromatin contacts with high accuracy genome-wide [98]

### Molecular Dynamics (MD) simulations

To derive a statistical ensemble of single-molecule model 3D structures of the *LTN1* locus, we performed extensive Molecular Dynamics (MD) simulations within the free available LAMMPS software [99] optimized for parallel computation [100,101]. In the model MD implementation, the polymer is a standard coarse-grained bead-spring chain [102] and the binders simple spherical particles. The motion of polymer beads and binders is determined by a Langevin equation with standard parameters (friction coefficient *ζ=*0.5 and temperature *T*=1, dimensionless units), numerically integrated via the Velocity-Verlet algorithm [103]. For simplicity, in our simulations, we set the diameter of both polymer beads and binders to *σ*=1, using it as the unit length of the model; similarly, we set the mass of beads and binders to be equal, taking it as the reference mass unit *m*=1 [102].

The physical interaction potentials are set as in classical polymer simulation studies [102]: excluded volume interactions between consecutive beads are modelled using a truncated, purely repulsive Lennard-Jones (LJ) potential; adjacent polymer beads are connected by FENE bonds with standard parameters (maximum length 1.6*σ* and spring constant 30K_B_T/*σ*^2^); attractive interactions between beads and cognate binders are described by a short-range, truncated LJ (cut-off distance 1.5*σ*). To test model robustness, we sampled a broad range of bead-binder affinity values in the weak biochemical energy scale, from 0 to 8K_B_T, and explored up to three orders of magnitude in binder concentration as detailed in previous studies [40,87]. For each set of parameters, we performed up to 3x10^2^ independent runs to ensure statistical strength.

The initial states of the MD simulations are open SAW configurations, located in a cubic simulation box of size 50*σ* with periodic boundary conditions to control finite size effects [104]. Binders are randomly injected in the simulation box, and then the system of beads and binders is equilibrated for 10^8^ MD time iteration steps. To monitor the system folding dynamics, we recorded the time track of the polymer gyration radius, *R*_g_. This function has a sharp drop at a characteristic time scale and then plateaus, marking the phase transition of the polymer from an initial free SAW chain to an equilibrium phase-separated globular conformation (**Figure 1b**). To give a sense of the chain compaction, the linear size of the polymer decreases by around 70% in the transition [96], resulting in an average *R*_g_ in the phase-separated state equal to 6.4*σ*.

The MD length scale (i.e., the bead diameter *σ*) can me mapped into physical units by using the relation [69]: *σ*=(*s/G*)*D*^1/3^, where *s* is the genomic content per bead (*s*=*L/N*, where *L*=2Mb is the genomic length of the locus and *N*=800 the number of beads), *G* the genome length (6Gb) and *D* the cell nucleus diameter (taken to be 10μm as an order of magnitude). These approximations yield *σ*=74.7 nm, in line with previous polymer physics studies [58,64]. As an additional check, single-cell microscopy data of the same *LTN1* genomic region in human IMR90 cells reported an average *R*_g_ of the locus equal to 464 nm [29]. This value, matched with that found in our simulations (6.4*σ*), returns *σ*=72.5 nm, remarkably close to our previous calculation. In the following, to set a reference value for *σ*, we take the average of those two independent estimates, i.e., *σ*=73.6 nm.

Finally, the simulation time, *τ*, can be converted into physical time via the formula: *τ =* 6πη*σ*^3^/(K_B_T), where η is the nucleoplasm viscosity. Given typical viscosity values in the range of a few fractions of poise [42,105], the model time scale is on the order of milliseconds, consistent with classic chromatin simulations [42].

## RESULTS

### Folding of the SBS model of the human *LTN1* gene locus

To comprehensively investigate the folding properties of our model at the ensemble-population level, we measured the average pairwise contact matrix in its phase-separated state. This was achieved by computing the mean of the model single-molecule contact maps, i.e., symmetric square matrices where each entry, *A*_ij_, is either 1 or 0, depending on whether the polymer sites *i* and *j* are in contact. A contact event is considered to occur if the spatial distance between the sites is below a typical distance threshold [42,83]. For our study, we explored thresholds within the range from 2 up to 5*σ*, corresponding to a spatial distance range of about 150-350 nm (see above), and they all provided analogous results. This range is consistent with established reference contact threshold values from microscopy studies [32].

To benchmark and validate the model output, we used publicly available in-situ Hi-C 2.0 data [93] of the *LTN1* locus in human WTC-11 cells from the 4DN Data Portal (dataset reference number 4DNESJ7S5NDJ) [94]. The experimental Hi-C contact map, which is binned at 25 kb resolution to match the model coarse-graining level, exhibits specific and non-random patterns of contacts, including the presence of TADs and sub-TAD domains, inter-TAD interactions, and long-range (>500kb) looping contacts, particularly around the *LTN1* gene (**Figure 2**, left panel).

**Figure 2.**
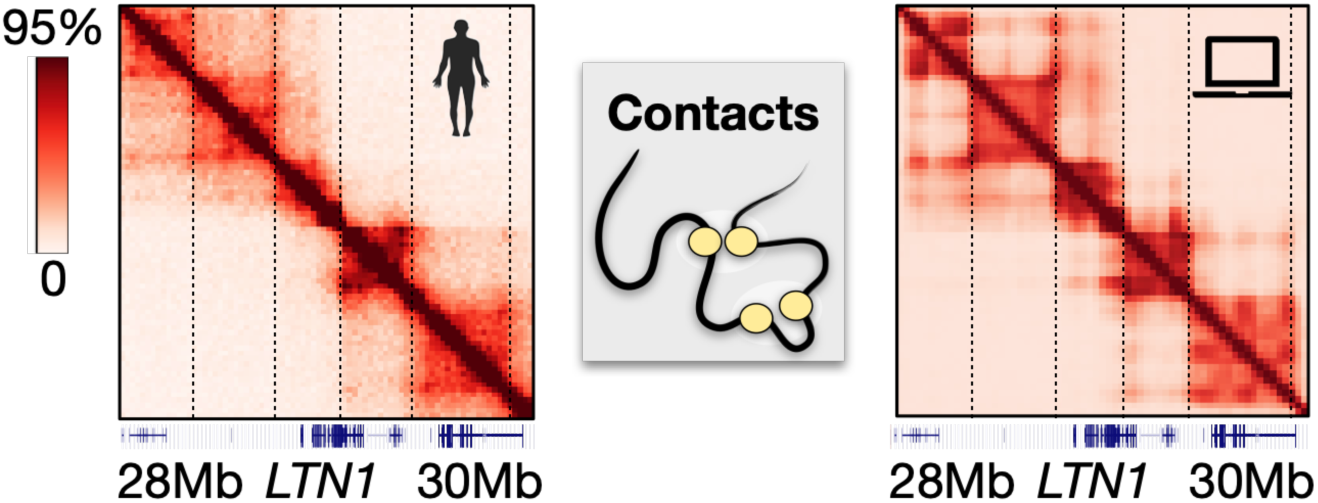
In-situ Hi-C 2.0 contact data of the studied 2Mb wide *LTN1* locus in WTC-11 cells (left) are consistently captured by the SBS polymer model (right). The high Pearson and genomic-distance corrected correlation values (respectively, r=0.90 and r’=0.59) indicate that the model accurately captures the overall structure of *LTN1* pairwise interactions.

Remarkably, the contact map generated by the model captures these observed experimental features, successfully recapitulating the overall structure of pairwise interactions (**Figure 2**, right). This agreement is quantitatively supported by the high Pearson correlation coefficient between the model and Hi-C contact matrices, r = 0.90. As an additional measure of similarity, we also computed the genomic-distance corrected Pearson correlation coefficient, r’, which averages out trivial genomic-proximity effects [83]. Despite the minimal complexity of the model, we find rʹ=0.59, which is a comparatively high correlation value considering that a randomly folded control chain would produce an r’ close to zero [106].

Thus, the ensemble of model conformations of the *LTN1* locus aligns well with Hi-C experimental data, demonstrating that our model quantitatively captures in-silico the structural features of the locus folding at the population-average level.

### Structural heterogeneity of *LTN1* single-molecule conformations in the model

Next, we aimed to study the folding dynamics of the locus as predicted by our simulations at the single-molecule level. By leveraging the 3D coordinates generated through the SBS model, we computed spatial distance maps for each phase-separated polymer conformation. These maps are symmetric square matrices of pairwise Euclidean distances between polymer sites across the locus. Our analysis reveals distinct single-molecule distance patterns, featuring TAD-like domains occurring at different genomic positions and long-range loop contacts spanning across TAD boundaries (**Figure 3a**). Consistent with previous studies [29,34], this variability underscores the highly dynamic nature of the 3D structures of single chromatin conformations. In the SBS model, such a structural heterogeneity arises, beyond stochastic thermal fluctuations, from the inherent folding degeneracy due to the specific, overlapping distribution of its binding sites [87] (see Figure 1a).

**Figure 3.**
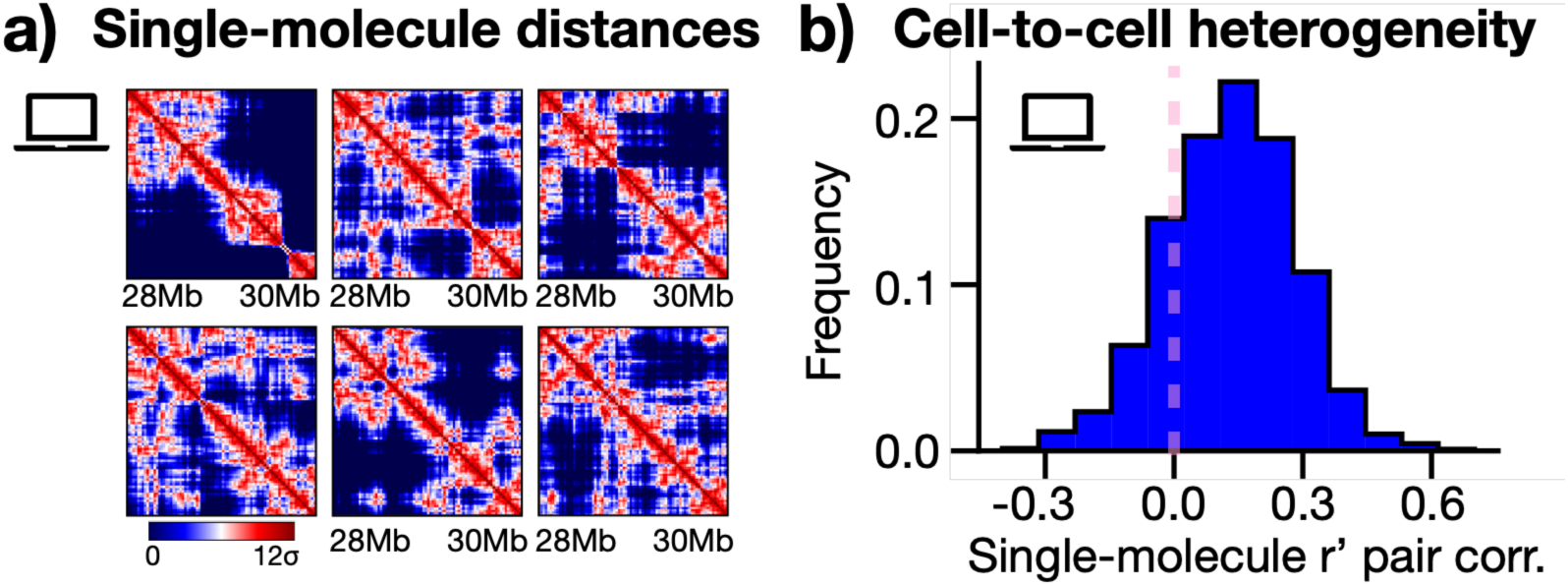
**a)** Representative examples of model-predicted phase-separated single-molecule distance matrices of the *LTN1* locus. The interaction patterns broadly differ across the ensemble of polymer configurations, as the system can fold in a variety of 3D architectures [87]. **b)** The structural heterogeneity of individual polymer structures is measured in the model by computing the r’ correlation between pairs of single-molecule distance matrices. The resulting distribution is broad (variance = 0.15) and has a non-zero average value (mean = 0.13), indicating that chromatin structures are highly variable from cell to cell, yet with a residual degree of structural correlation.

To quantify the extent of structural variability among model single molecules, we measured the r’ pairwise correlations between their distance matrices. In the absence of correlation between matrices, we would expect an r’ distribution centered around zero. Conversely, perfectly correlated matrices would yield an r’ distribution peaked at 1. Interestingly, our calculations return a unimodal distribution with a mean value of 0.13 and a variance of 0.15 (**Figure 3b**): that suggests a substantial heterogeneity of polymer 3D structures that, though, retain a residual degree of structural correlation (as indicated by the non-zero r’ average value). These results are in line with recent microscopy studies that reported a strong cell-to-cell variability of individual chromatin conformations at the sub-megabase scale [29,30,33], revealing, nevertheless, sub-clusters of structures with correlated behaviors [32].

Summarizing, the model highlights a broad distribution of single molecule 3D structures at the human *LTN1* locus, providing quantitative predictions (e.g., single-molecule spatial distances and all-against-all pair correlations) which can be directly tested by independent single-cell super-resolution microscopy approaches. Such experimental validation would not only assess the predictive power of the model but also deepen our understanding of the intrinsic structural diversity and dynamic behavior of genomic loci at the single-molecule level.

### Shape and size of 3D model single molecules

To further characterize the *LTN1* single-molecule structures predicted by our chromatin model, we calculated their shape and volume. Specifically, for each model conformation in the phase-separated state, we computed the inertia tensor, defined as: 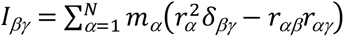, where *N* is the number of monomers along the polymer chain, *m_α_* the mass of the *α-*th monomer, *r_αβ_* its *β*-th spatial coordinate and *β,γ* are spatial component indexes equal to {0,1,2}. The eigenvalues of this tensor, corresponding to the system principal moments of inertia *I_a_*, *I_b_* and *I_c_*, can be related through standard textbook formulas to the semi-axes (*a*, *b*, *c*) of a triaxial ellipsoid enclosing the volume contour of a given conformation: 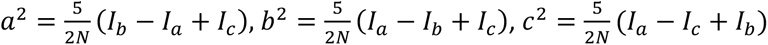.

In the case of a perfectly spherical conformation, the three eigenvalues are equal and that implies *a* = *b* = *c*. However, in the case of the *LTN1* locus, our model predicts substantial deviations of individual chromatin conformations from a spherical topology, as they are found to have a prolate shape with *a* > *b* ≈ *c* (**Figure 4a**). Indeed, while the distribution of the ratios *b*/*c* has an average value of 1.0 (orange histogram in the figure), the ratios *a*/*c* are centered around 2.0 (blue histogram); moreover, statistical analysis indicates that these two distributions are significantly distinguishable from each other (two-sided Mann-Whitney p-value <0.001).

**Figure 4.**
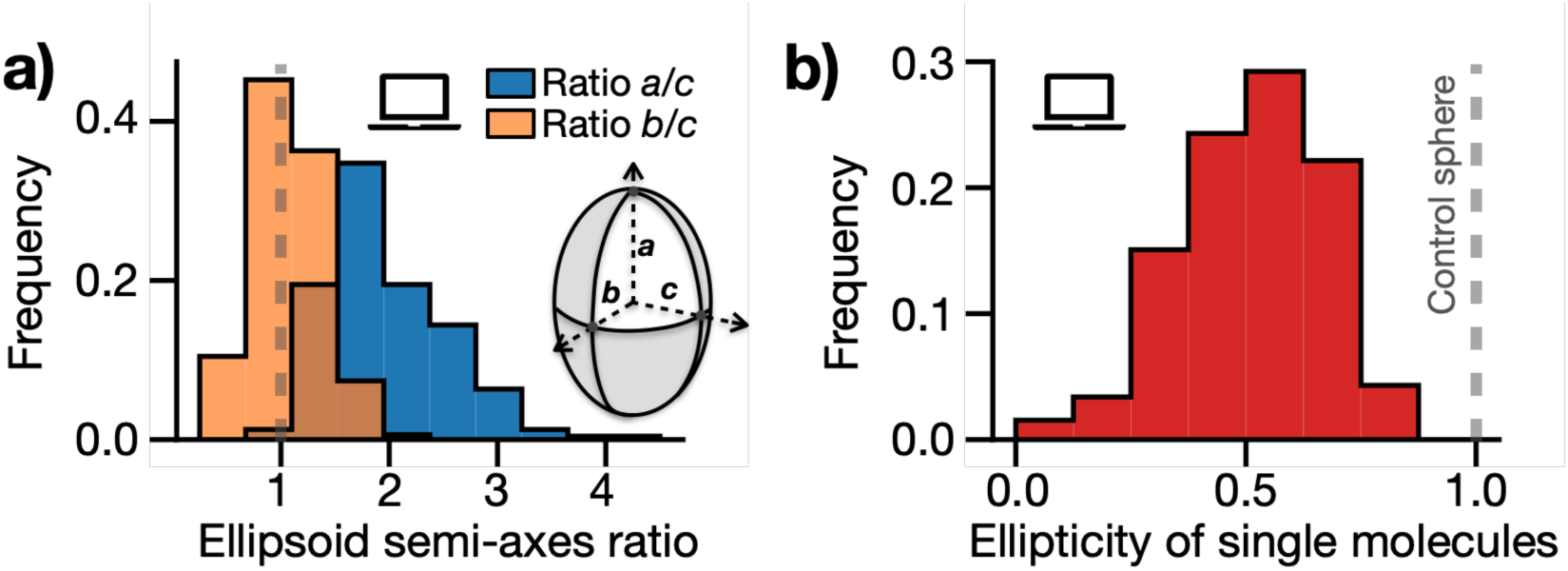
**a)** Distribution of ellipsoid semi-axes ratios, *a*/*c* (histogram in blue) and *b*/*c* (in orange), computed from the inertia tensor of single-molecule polymer structures. The black dashed line in the figure represents the expected value in the case of perfectly spherical conformations (*a*=*b*=*c*). Interestingly, model-predicted 3D structures of the *LTN1* locus appear prolate, as we find *a* > *b* ≈ *c*. The two distributions *a*/*c* and *b*/*c* are statistically different from each other (two-sided Mann-Whitney p-value <0.001). **b)** The ellipticity of model single molecules, calculated from the eigenvalues of their gyration tensor [107], exhibits a broad distribution (variance=0.16) with an average value of 0.51, indicating a significant structural variability and a tendency towards a prolate shape.

By using the inertia tensor, we also computed the volume *V* of single-molecule structures via the formula *V*=4/3π*abc*, which provides an average volume estimation of the entire locus of 1500*σ*^3^. To give a sense of the physical length scales, by taking *σ* = 73.6 nm (see above), the model predicts that the average volume of the locus is thus approximately 0.59 μm³. We also checked an alternative volume calculation using the formula *V*=4/3π*R*_g3_, which returned a mean *LTN1* volume equal to 1300*σ*^3^ (i.e., 0.52 μm³) comparable to our previous estimate. Overall, these findings are consistent with typical volume size estimates of Mb-wide genomic loci, falling in the range of fractions of a few μm^3^, as measured, e.g., by super-resolution microscopy experiments [29].

As an additional test on the shape of model-predicted conformations, we computed the polymer gyration tensor, defined as [107]: 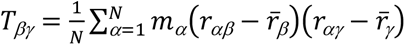, where *r_βγ_* (*r_α_γ*) is the *β*-th (*γ*-th) component of the position vector of the *α*-th monomer and *r̄_β_* (*r̄_γ_*) the component of mass center of the polymer chain along the *β*-th (*γ*-th) direction. By diagonalizing the tensor, we derived its three eigenvalues, λ_1_ ζ λ_2_ ζ λ_3_, from which we calculated the ellipticity, ε, of the model single molecules, defined as ε=2λ_3_/(λ_1_+λ_2_). For spherical configurations ε=1, as the three eigenvalues are expected to be equal [107]. Consistent with the observed high structural variability of the *LTN1* polymer conformations, we find that distribution of single-molecule ellipticity values is broad (variance = 0.16), yet it has an average of 0.51 well below that of a spherical control (**Figure 4b**), highlighting the prolate nature of the single-molecule structures predicted by the model.

Taken together, these results indicate that the *LTN1* locus in human WTC-11 cells significantly deviates from a simple spherical geometry, predominantly displaying a prolate shape with substantial single-molecule structural variability, as predicted by our chromatin model and tested through multiple, independent shape and volume metrics.

## DISCUSSION

In this work, we combined polymer physics models of chromatin and computer simulations to comprehensively analyze the folding properties of the *LTN1* gene locus in a human induced pluripotent stem cell line. By computing the average pairwise contact matrix of the model, we demonstrated that its predicted patterns are consistent with available in-situ Hi-C 2.0 experiments [93,94], capturing with high accuracy the complex architecture of TADs, sub-TADs, and long-range looping interactions observed at the gene locus.

Beyond population averages, our model reveals significant structural heterogeneity among individual chromatin 3D structures. This variability is evidenced, for instance, by distinct folding patterns in single-molecule spatial distance matrices and the broad distribution of their pairwise correlation values. Our results suggest that while the *LTN1* locus exhibits a substantial range of conformations, it also maintains a degree of structural correlation, consistent with observations from recent microscopy studies [29,32].

Furthermore, our calculations of the inertia and gyration tensors provide quantitative insights into the shape and volume of the locus conformations. The model predicts a prolate shape of single molecules, with an average volume estimate in line with super-resolution imaging data [29]. It also provides additional testable predictions, such as the distribution of single-molecule ellipticity values, which can be directly validated by advanced microscopy techniques. Albeit simplified, the model is robust as it is dictated by thermodynamics, and phase transitions and complex emergent behaviors are common to biological and systems of soft-matter physics [108–119].

These findings have significant implications beyond structural characterization. Understanding the variability and dynamics of genomic loci at the single-molecule level is crucial for elucidating the principles of gene regulation, chromatin organization, and cellular function. The observed heterogeneity in 3D structures, indeed, could influence gene expression patterns, regulatory element interactions, and overall genomic stability.

The ability to predict and verify the structural configurations of genomic loci through principled computational models enhances our capacity to interpret how physical changes in individual chromatin structures can lead to functional outcomes, such as differential gene expression, cellular differentiation and genetic-disease-associated phenotypes. That provides a robust basis for experimental validation at the single-cell level and a deeper understanding of the dynamic and heterogeneous nature of chromatin 3D architectures.

## AUTHOR CONTRIBUTIONS

Conceptualization, M.C. and M.N.; writing—original draft preparation, M.C. and A.A.; formal analysis, M.C., A.A. and A.E.; data curation and production, L.Y., J.H.G., K.M.P. and J.D.; visualization, M.C., A.A, A.E., F.V., A.F. and F.D.P.; supervision, M.C and M.N. All authors have read and agreed to the published version of the manuscript.

## DATA AVAILABILITY

The data that support the findings of this study are available from the corresponding author upon reasonable request. Hi-C 2.0 data used in this study are available at the 4DN Data Portal [94] (dataset reference number 4DNESJ7S5NDJ).

## ACKNOWLEDGMENTS

We acknowledge computer resources from INFN, CINECA, ENEA CRESCO/ENEAGRID [120] and *Scope*/*ReCAS*/*Ibisco* at the University of Naples. J.D. is an investigator of the Howard Hughes Medical Institute.

## REFERENCES

1. Lieberman-Aiden, E.; van Berkum, N.L.; Williams, L.; Imakaev, M.; Ragoczy, T.; Telling, A.; Amit, I.; Lajoie, B.R.; Sabo, P.J.; Dorschner, M.O.;, et al. Comprehensive Mapping of Long-Range Interactions Reveals Folding Principles of the Human Genome. Science (80-.). 2009, 326, 289–293, doi:10.1126/science.1181369.

2. Belton, J.M.; McCord, R.P.; Gibcus, J.H.; Naumova, N.; Zhan, Y.; Dekker, J. Hi-C: A Comprehensive Technique to Capture the Conformation of Genomes. Methods 2012, 58, doi:10.1016/j.ymeth.2012.05.001.

3. Beagrie, R.A.; Scialdone, A.; Schueler, M.; Kraemer, D.C.A.; Chotalia, M.; Xie, S.Q.; Barbieri, M.; De Santiago, I.; Lavitas, L.M.; Branco, M.R.;, et al. Complex Multi-Enhancer Contacts Captured by Genome Architecture Mapping. Nature 2017, 543, 519–524, doi:10.1038/nature21411.

4. Winick-Ng, W.; Kukalev, A.; Harabula, I.; Zea-Redondo, L.; Szabó, D.; Meijer, M.; Serebreni, L.; Zhang, Y.; Bianco, S.; Chiariello, A.M.;, et al. Cell-Type Specialization Is Encoded by Specific Chromatin Topologies. Nature 2021, 599, 684–691, doi:10.1038/s41586-021-04081-2.

5. Quinodoz, S.A.; Ollikainen, N.; Tabak, B.; Palla, A.; Schmidt, J.M.; Detmar, E.; Lai, M.M.; Shishkin, A.A.; Bhat, P.; Takei, Y.;, et al. Higher-Order Inter-Chromosomal Hubs Shape 3D Genome Organization in the Nucleus. Cell 2018, 174, 744–757.e24, doi:10.1016/j.cell.2018.05.024.

6. Quinodoz, S.A.; Bhat, P.; Chovanec, P.; Jachowicz, J.W.; Ollikainen, N.; Detmar, E.; Soehalim, E.; Guttman, M. SPRITE: A Genome-Wide Method for Mapping Higher-Order 3D Interactions in the Nucleus Using Combinatorial Split-and-Pool Barcoding. Nat. Protoc. 2022, 17, 36–75, doi:10.1038/s41596-021-00633-y.

7. Bickmore, W.A.; van Steensel, B. Genome Architecture: Domain Organization of Interphase Chromosomes. Cell 2013, 152, 1270–1284, doi:10.1016/j.cell.2013.02.001.

8. Dekker, J.; Mirny, L. The 3D Genome as Moderator of Chromosomal Communication. Cell 2016, 164, 1110–1121, doi:10.1016/j.cell.2016.02.007.

9. Dixon, J.R.; Gorkin, D.U.; Ren, B. Chromatin Domains: The Unit of Chromosome Organization. Mol. Cell 2016, 62, 668–680, doi:10.1016/j.molcel.2016.05.018.

10. Spielmann, M.; Lupiáñez, D.G.; Mundlos, S. Structural Variation in the 3D Genome. Nat. Rev. Genet. 2018, 19, 453–467, doi:10.1038/s41576-018-0007-0.

11. Akgol Oksuz, B.; Yang, L.; Abraham, S.; Venev, S. V.; Krietenstein, N.; Parsi, K.M.; Ozadam, H.; Oomen, M.E.; Nand, A.; Mao, H.;, et al. Systematic Evaluation of Chromosome Conformation Capture Assays. Nat. Methods 2021, 18, doi:10.1038/s41592-021-01248-7.

12. Rao, S.S.P.; Huntley, M.H.; Durand, N.C.; Stamenova, E.K.; Bochkov, I.D.; Robinson, J.T.; Sanborn, A.L.; Machol, I.; Omer, A.D.; Lander, E.S.;, et al. A 3D Map of the Human Genome at Kilobase Resolution Reveals Principles of Chromatin Looping. Cell 2014, 159, 1665–1680, doi:10.1016/j.cell.2014.11.021.

13. Nora, E.P.; Lajoie, B.R.; Schulz, E.G.; Giorgetti, L.; Okamoto, I.; Servant, N.; Piolot, T.; van Berkum, N.L.; Meisig, J.; Sedat, J.;, et al. Spatial Partitioning of the Regulatory Landscape of the X-Inactivation Centre. Nature 2012, 485, 381–385, doi:10.1038/nature11049.

14. Dixon, J.R.; Selvaraj, S.; Yue, F.; Kim, A.; Li, Y.; Shen, Y.; Hu, M.; Liu, J.S.; Ren, B. Topological Domains in Mammalian Genomes Identified by Analysis of Chromatin Interactions. Nature 2012, 485, 376–380, doi:10.1038/nature11082.

15. Fraser, J.; Ferrai, C.; Chiariello, A.M.; Schueler, M.; Rito, T.; Laudanno, G.; Barbieri, M.; Moore, B.L.; Kraemer, D.C.; Aitken, S.;, et al. Hierarchical Folding and Reorganization of Chromosomes Are Linked to Transcriptional Changes in Cellular Differentiation. Mol. Syst. Biol. 2015, doi:10.15252/msb.20156492.

16. van Steensel, B.; Belmont, A.S. Lamina-Associated Domains: Links with Chromosome Architecture, Heterochromatin, and Gene Repression. Cell 2017, 169, 780–791, doi:10.1016/j.cell.2017.04.022.

17. Cremer, T.; Cremer, C. Chromosome Territories, Nuclear Architecture and Gene Regulation in Mammalian Cells. Nat. Rev. Genet. 2001, 2, 292–301, doi:10.1038/35066075.

18. Sexton, T.; Cavalli, G. The Role of Chromosome Domains in Shaping the Functional Genome. Cell 2015, 160, 1049–1059, doi:10.1016/j.cell.2015.02.040.

19. Schoenfelder, S.; Fraser, P. Long-Range Enhancer–Promoter Contacts in Gene Expression Control. Nat. Rev. Genet. 2019, 20.

20. Kubo, N.; Ishii, H.; Xiong, X.; Bianco, S.; Meitinger, F.; Hu, R.; Hocker, J.D.; Conte, M.; Gorkin, D.; Yu, M.;, et al. Promoter-Proximal CTCF Binding Promotes Distal Enhancer-Dependent Gene Activation. Nat. Struct. Mol. Biol. 2021, 28, 152–161, doi:10.1038/s41594-020-00539-5.

21. Huang, H.; Zhu, Q.; Jussila, A.; Han, Y.; Bintu, B.; Kern, C.; Conte, M.; Zhang, Y.; Bianco, S.; Chiariello, A.M.;, et al. CTCF Mediates Dosage- and Sequence-Context-Dependent Transcriptional Insulation by Forming Local Chromatin Domains. Nat. Genet. 2021, 53, 1064– 1074, doi:10.1038/s41588-021-00863-6.

22. Willemin, A.; Szabó, D.; Pombo, A. Epigenetic Regulatory Layers in the 3D Nucleus. Mol. Cell 2024, 84.

23. Salamon, I.; Serio, S.; Bianco, S.; Pagiatakis, C.; Crasto, S.; Chiariello, A.M.; Conte, M.; Cattaneo, P.; Fiorillo, L.; Felicetta, A.;, et al. Divergent Transcription of the Nkx2-5 Locus Generates Two Enhancer RNAs with Opposing Functions. iScience 2020, 23.

24. Lupiáñez, D.G.; Kraft, K.; Heinrich, V.; Krawitz, P.; Brancati, F.; Klopocki, E.; Horn, D.; Kayserili, H.; Opitz, J.M.; Laxova, R.;, et al. Disruptions of Topological Chromatin Domains Cause Pathogenic Rewiring of Gene-Enhancer Interactions. Cell 2015, 161, doi:10.1016/j.cell.2015.04.004.

25. Lupiáñez, D.G.; Spielmann, M.; Mundlos, S. Breaking TADs: How Alterations of Chromatin Domains Result in Disease. Trends Genet. 2016, 32, 225–237, doi:10.1016/j.tig.2016.01.003.

26. Hnisz, D.; Weintraub, A.S.; Day, D.S.; Valton, A.-L.; Bak, R.O.; Li, C.H.; Goldmann, J.; Lajoie, B.R.; Fan, Z.P.; Sigova, A.A.;, et al. Activation of Proto-Oncogenes by Disruption of Chromosome Neighborhoods. Science (80-.). 2016, 351, 1454–1458, doi:10.1126/science.aad9024.

27. Franke, M.; Ibrahim, D.M.; Andrey, G.; Schwarzer, W.; Heinrich, V.; Schöpflin, R.; Kraft, K.; Kempfer, R.; Jerković, I.; Chan, W.-L.;, et al. Formation of New Chromatin Domains Determines Pathogenicity of Genomic Duplications. Nature 2016, 538, 265–269, doi:10.1038/nature19800.

28. Finn, E.H.; Misteli, T. Molecular Basis and Biological Function of Variability in Spatial Genome Organization. Science (80-.). 2019, 365, eaaw9498, doi:10.1126/science.aaw9498.

29. Bintu, B.; Mateo, L.J.; Su, J.-H.; Sinnott-Armstrong, N.A.; Parker, M.; Kinrot, S.; Yamaya, K.; Boettiger, A.N.; Zhuang, X. Super-Resolution Chromatin Tracing Reveals Domains and Cooperative Interactions in Single Cells. Science (80-.). 2018, 362, eaau1783, doi:10.1126/science.aau1783.

30. Su, J.-H.; Zheng, P.; Kinrot, S.S.; Bintu, B.; Zhuang, X. Genome-Scale Imaging of the 3D Organization and Transcriptional Activity of Chromatin. Cell 2020, 182, 1641–1659.e26, doi:10.1016/j.cell.2020.07.032.

31. Takei, Y.; Yun, J.; Zheng, S.; Ollikainen, N.; Pierson, N.; White, J.; Shah, S.; Thomassie, J.; Suo, S.; Eng, C.-H.L.;, et al. Integrated Spatial Genomics Reveals Global Architecture of Single Nuclei. Nature 2021, 590, 344–350, doi:10.1038/s41586-020-03126-2.

32. Takei, Y.; Zheng, S.; Yun, J.; Shah, S.; Pierson, N.; White, J.; Schindler, S.; Tischbirek, C.H.; Yuan, G.-C.; Cai, L. Single-Cell Nuclear Architecture across Cell Types in the Mouse Brain. Science (80-.). 2021, 374, 586–594, doi:10.1126/science.abj1966.

33. Szabo, Q.; Donjon, A.; Jerković, I.; Papadopoulos, G.L.; Cheutin, T.; Bonev, B.; Nora, E.P.; Bruneau, B.G.; Bantignies, F.; Cavalli, G. Regulation of Single-Cell Genome Organization into TADs and Chromatin Nanodomains. Nat. Genet. 2020, 52, 1151–1157, doi:10.1038/s41588-020-00716-8.

34. Finn, E.H.; Pegoraro, G.; Brandão, H.B.; Valton, A.-L.; Oomen, M.E.; Dekker, J.; Mirny, L.; Misteli, T. Extensive Heterogeneity and Intrinsic Variation in Spatial Genome Organization. Cell 2019, 176, 1502–1515.e10, doi:10.1016/j.cell.2019.01.020.

35. Boettiger, A.N.; Bintu, B.; Moffitt, J.R.; Wang, S.; Beliveau, B.J.; Fudenberg, G.; Imakaev, M.; Mirny, L.A.; Wu, C.; Zhuang, X. Super-Resolution Imaging Reveals Distinct Chromatin Folding for Different Epigenetic States. Nature 2016, 529, 418–422, doi:10.1038/nature16496.

36. Götz, M.; Messina, O.; Espinola, S.; Fiche, J.-B.; Nollmann, M. Multiple Parameters Shape the 3D Chromatin Structure of Single Nuclei at the Doc Locus in Drosophila. Nat. Commun. 2022, 13, 5375, doi:10.1038/s41467-022-32973-y.

37. Jerković, I.; Cavalli, G. Understanding 3D Genome Organization by Multidisciplinary Methods. Nat. Rev. Mol. Cell Biol. 2021, 22, 511–528, doi:10.1038/s41580-021-00362-w.

38. Li, Q.; Tjong, H.; Li, X.; Gong, K.; Zhou, X.J.; Chiolo, I.; Alber, F. The Three-Dimensional Genome Organization of Drosophila Melanogaster through Data Integration. Genome Biol. 2017, 18, 145, doi:10.1186/s13059-017-1264-5.

39. Tjong, H.; Li, W.; Kalhor, R.; Dai, C.; Hao, S.; Gong, K.; Zhou, Y.; Li, H.; Zhou, X.J.; Le Gros, M.A.;, et al. Population-Based 3D Genome Structure Analysis Reveals Driving Forces in Spatial Genome Organization. Proc. Natl. Acad. Sci. U. S. A. 2016, doi:10.1073/pnas.1512577113.

40. Chiariello, A.M.; Annunziatella, C.; Bianco, S.; Esposito, A.; Nicodemi, M. Polymer Physics of Chromosome Large-Scale 3D Organisation. Sci. Rep. 2016, 6, 29775, doi:10.1038/srep29775.

41. Chiariello, A.M.; Bianco, S.; Esposito, A.; Fiorillo, L.; Conte, M.; Irani, E.; Musella, F.; Abraham, A.; Prisco, A.; Nicodemi, M. Physical Mechanisms of Chromatin Spatial Organization. FEBS J. 2022, 289, 1180–1190, doi:10.1111/febs.15762.

42. Brackley, C.A.; Taylor, S.; Papantonis, A.; Cook, P.R.; Marenduzzo, D. Nonspecific Bridging-Induced Attraction Drives Clustering of DNA-Binding Proteins and Genome Organization. Proc. Natl. Acad. Sci. 2013, 110, E3605–E3611, doi:10.1073/pnas.1302950110.

43. Brackley, C.A.; Liebchen, B.; Michieletto, D.; Mouvet, F.; Cook, P.R.; Marenduzzo, D. Ephemeral Protein Binding to DNA Shapes Stable Nuclear Bodies and Chromatin Domains. Biophys. J. 2017, 112, 1085–1093, doi:10.1016/j.bpj.2017.01.025.

44. Nir, G.; Farabella, I.; Pérez Estrada, C.; Ebeling, C.G.; Beliveau, B.J.; Sasaki, H.M.; Lee, S.H.; Nguyen, S.C.; McCole, R.B.; Chattoraj, S.;, et al. Walking along Chromosomes with Super-Resolution Imaging, Contact Maps, and Integrative Modeling. PLoS Genet. 2018, 14, e1007872, doi:10.1371/journal.pgen.1007872.

45. Di Pierro, M.; Zhang, B.; Aiden, E.L.; Wolynes, P.G.; Onuchic, J.N. Transferable Model for Chromosome Architecture. Proc. Natl. Acad. Sci. 2016, 113, 12168–12173, doi:10.1073/pnas.1613607113.

46. Rosa, A.; Everaers, R. Structure and Dynamics of Interphase Chromosomes. PLoS Comput. Biol. 2008, 4, e1000153, doi:10.1371/journal.pcbi.1000153.

47. Esposito, A.; Bianco, S.; Fiorillo, L.; Conte, M.; Abraham, A.; Musella, F.; Nicodemi, M.; Prisco, A.; Chiariello, A.M. Polymer Models Are a Versatile Tool to Study Chromatin 3D Organization. Biochem. Soc. Trans. 2021, 49, 1675–1684, doi:10.1042/BST20201004.

48. Bianco, S.; Annunziatella, C.; Andrey, G.; Chiariello, A.M.; Esposito, A.; Fiorillo, L.; Prisco, A.; Conte, M.; Campanile, R.; Nicodemi, M. Modeling Single-Molecule Conformations of the HoxD Region in Mouse Embryonic Stem and Cortical Neuronal Cells. Cell Rep. 2019, 28, 1574–1583.e4, doi:10.1016/j.celrep.2019.07.013.

49. Bianco, S.; Chiariello, A.M.; Conte, M.; Esposito, A.; Fiorillo, L.; Musella, F.; Nicodemi, M. Computational Approaches from Polymer Physics to Investigate Chromatin Folding. Curr. Opin. Cell Biol. 2020, 64, 10–17, doi:10.1016/j.ceb.2020.01.002.

50. Serra, F.; Baù, D.; Goodstadt, M.; Castillo, D.; Filion, G.J.; Marti-Renom, M.A. Automatic Analysis and 3D-Modelling of Hi-C Data Using TADbit Reveals Structural Features of the Fly Chromatin Colors. PLOS Comput. Biol. 2017, 13, e1005665, doi:10.1371/journal.pcbi.1005665.

51. Neguembor, M.V.; Arcon, J.P.; Buitrago, D.; Lema, R.; Walther, J.; Garate, X.; Martin, L.; Romero, P.; AlHaj Abed, J.; Gut, M.;, et al. MiOS, an Integrated Imaging and Computational Strategy to Model Gene Folding with Nucleosome Resolution. Nat. Struct. Mol. Biol. 2022, 29, 1011–1023, doi:10.1038/s41594-022-00839-y.

52. Racko, D.; Benedetti, F.; Dorier, J.; Stasiak, A. Transcription-Induced Supercoiling as the Driving Force of Chromatin Loop Extrusion during Formation of TADs in Interphase Chromosomes. Nucleic Acids Res. 2018, 46, 1648–1660, doi:10.1093/nar/gkx1123.

53. Sanborn, A.L.; Rao, S.S.P.; Huang, S.C.; Durand, N.C.; Huntley, M.H.; Jewett, A.I.; Bochkov, I.D.; Chinnappan, D.; Cutkosky, A.; Li, J.;, et al. Chromatin Extrusion Explains Key Features of Loop and Domain Formation in Wild-Type and Engineered Genomes. Proc. Natl. Acad. Sci. U. S. A. 2015, 112, E6456–E6465, doi:10.1073/pnas.1518552112.

54. Fudenberg, G.; Imakaev, M.; Lu, C.; Goloborodko, A.; Abdennur, N.; Mirny, L.A. Formation of Chromosomal Domains by Loop Extrusion. Cell Rep. 2016, 15, 2038–2049, doi:10.1016/j.celrep.2016.04.085.

55. Yildirim, A.; Boninsegna, L.; Zhan, Y.; Alber, F. Uncovering the Principles of Genome Folding by 3D Chromatin Modeling. Cold Spring Harb. Perspect. Biol. 2022, 14, a039693, doi:10.1101/cshperspect.a039693.

56. Boninsegna, L.; Yildirim, A.; Polles, G.; Zhan, Y.; Quinodoz, S.A.; Finn, E.H.; Guttman, M.; Zhou, X.J.; Alber, F. Integrative Genome Modeling Platform Reveals Essentiality of Rare Contact Events in 3D Genome Organizations. Nat. Methods 2022, 19, 938–949, doi:10.1038/s41592-022-01527-x.

57. Esposito, A.; Abraham, A.; Conte, M.; Vercellone, F.; Prisco, A.; Bianco, S.; Chiariello, A.M. The Physics of DNA Folding: Polymer Models and Phase-Separation. Polymers (Basel). 2022, 14, 1918, doi:10.3390/polym14091918.

58. Conte, M.; Irani, E.; Chiariello, A.M.; Abraham, A.; Bianco, S.; Esposito, A.; Nicodemi, M. Loop-Extrusion and Polymer Phase-Separation Can Co-Exist at the Single-Molecule Level to Shape Chromatin Folding. Nat. Commun. 2022, 13, 4070.

59. Fiorillo, L.; Bianco, S.; Esposito, A.; Conte, M.; Sciarretta, R.; Musella, F.; Chiariello, A.M. A Modern Challenge of Polymer Physics: Novel Ways to Study, Interpret, and Reconstruct Chromatin Structure. WIREs Comput. Mol. Sci. 2020, 10, doi:10.1002/wcms.1454.

60. Esposito, A.; Chiariello, A.M.; Conte, M.; Fiorillo, L.; Musella, F.; Sciarretta, R.; Bianco, S. Higher-Order Chromosome Structures Investigated by Polymer Physics in Cellular Morphogenesis and Differentiation. J. Mol. Biol. 2020.

61. Lin, D.; Bonora, G.; Yardimci, G.G.; Noble, W.S. Computational Methods for Analyzing and Modeling Genome Structure and Organization. Wiley Interdiscip. Rev. Syst. Biol. Med. 2018, 11, e1435, doi:10.1002/wsbm.1435.

62. Zhang, B.; Wolynes, P.G. Topology, Structures, and Energy Landscapes of Human Chromosomes. Proc. Natl. Acad. Sci. 2015, 112, 6062–6067, doi:10.1073/pnas.1506257112.

63. Shi, G.; Thirumalai, D. From Hi-C Contact Map to Three-Dimensional Organization of Interphase Human Chromosomes. Phys. Rev. X 2021, 11, 011051, doi:10.1103/PhysRevX.11.011051.

64. Shi, G.; Liu, L.; Hyeon, C.; Thirumalai, D. Interphase Human Chromosome Exhibits out of Equilibrium Glassy Dynamics. Nat. Commun. 2018, 9, 3161, doi:10.1038/s41467-018-05606-6.

65. Brackley, C.A.; Johnson, J.; Michieletto, D.; Morozov, A.N.; Nicodemi, M.; Cook, P.R.; Marenduzzo, D. Nonequilibrium Chromosome Looping via Molecular Slip Links. Phys. Rev. Lett. 2017, 119, 138101, doi:10.1103/PhysRevLett.119.138101.

66. Plewczynski, D.; Kadlof, M. Computational Modelling of Three-Dimensional Genome Structure. Methods 2020, 181–182, 1–4, doi:10.1016/j.ymeth.2020.09.013.

67. Banigan, E.J.; Mirny, L.A. Loop Extrusion: Theory Meets Single-Molecule Experiments. Curr. Opin. Cell Biol. 2020, 64, 124–138, doi:10.1016/j.ceb.2020.04.011.

68. Nicodemi, M.; Prisco, A. Thermodynamic Pathways to Genome Spatial Organization in the Cell Nucleus. Biophys. J. 2009, 96, 2168–2177, doi:10.1016/j.bpj.2008.12.3919.

69. Barbieri, M.; Chotalia, M.; Fraser, J.; Lavitas, L.-M.; Dostie, J.; Pombo, A.; Nicodemi, M. Complexity of Chromatin Folding Is Captured by the Strings and Binders Switch Model. Proc. Natl. Acad. Sci. 2012, 109, 16173–16178, doi:10.1073/pnas.1204799109.

70. Crippa, M.; Zhan, Y.; Tiana, G. Effective Model of Loop Extrusion Predicts Chromosomal Domains. Phys. Rev. E 2020, 102, 032414, doi:10.1103/PhysRevE.102.032414.

71. Conte, M.; Esposito, A.; Vercellone, F.; Abraham, A.; Bianco, S. Unveiling the Machinery behind Chromosome Folding by Polymer Physics Modeling. Int. J. Mol. Sci. 2023, 24, 3660, doi:10.3390/ijms24043660.

72. Di Stefano, M.; Paulsen, J.; Lien, T.G.; Hovig, E.; Micheletti, C. Hi-C-Constrained Physical Models of Human Chromosomes Recover Functionally-Related Properties of Genome Organization. Sci. Rep. 2016, 6, 35985, doi:10.1038/srep35985.

73. Buckle, A.; Brackley, C.A.; Boyle, S.; Marenduzzo, D.; Gilbert, N. Polymer Simulations of Heteromorphic Chromatin Predict the 3D Folding of Complex Genomic Loci. Mol. Cell 2018, 72, 786–797.e11, doi:10.1016/j.molcel.2018.09.016.

74. Brackley, C.A.; Johnson, J.; Michieletto, D.; Morozov, A.N.; Nicodemi, M.; Cook, P.R.; Marenduzzo, D. Extrusion without a Motor: A New Take on the Loop Extrusion Model of Genome Organization. Nucleus 2018, 9, doi:10.1080/19491034.2017.1421825.

75. Conte, M.; Esposito, A.; Fiorillo, L.; Annunziatella, C.; Corrado, A.; Musella, F.; Sciarretta, R.; Chiariello, A.M.; Bianco, S. Hybrid Machine Learning and Polymer Physics Approach to Investigate 3D Chromatin Structure. In Proceedings of the Lecture Notes in Computer Science (including subseries Lecture Notes in Artificial Intelligence and Lecture Notes in Bioinformatics); 2020; Vol. 11997 LNCS.

76. Belokopytova, P.; Viesná, E.; Chiliński, M.; Qi, Y.; Salari, H.; Di Stefano, M.; Esposito, A.; Conte, M.; Chiariello, A.M.; Teif, V.B.;, et al. 3DGenBench: A Web-Server to Benchmark Computational Models for 3D Genomics. Nucleic Acids Res. 2022, 50, W4–W12, doi:10.1093/nar/gkac396.

77. Salari, H.; Di Stefano, M.; Jost, D. Spatial Organization of Chromosomes Leads to Heterogeneous Chromatin Motion and Drives the Liquid- or Gel-like Dynamical Behavior of Chromatin. Genome Res. 2022, 32, 28–43, doi:10.1101/gr.275827.121.

78. Jost, D.; Carrivain, P.; Cavalli, G.; Vaillant, C. Modeling Epigenome Folding: Formation and Dynamics of Topologically Associated Chromatin Domains. Nucleic Acids Res. 2014, 42, 9553– 9561, doi:10.1093/nar/gku698.

79. Tortora, M.M.; Salari, H.; Jost, D. Chromosome Dynamics during Interphase: A Biophysical Perspective. Curr. Opin. Genet. Dev. 2020, 61, 37–43, doi:10.1016/j.gde.2020.03.001.

80. Bohn, M.; Heermann, D.W. Diffusion-Driven Looping Provides a Consistent Framework for Chromatin Organization. PLoS One 2010, 5, e12218, doi:10.1371/journal.pone.0012218.

81. Conte, M.; Fiorillo, L.; Bianco, S.; Chiariello, A.M.; Esposito, A.; Musella, F.; Flora, F.; Abraham, A.; Nicodemi, M. A Polymer Physics Model to Dissect Genome Organization in Healthy and Pathological Phenotypes. In Methods in Molecular Biology; 2022; Vol. 2301, pp. 307–316.

82. Fudenberg, G.; Kelley, D.R.; Pollard, K.S. Predicting 3D Genome Folding from DNA Sequence with Akita. Nat. Methods 2020, 17, 1111–1117, doi:10.1038/s41592-020-0958-x.

83. Bianco, S.; Lupiáñez, D.G.; Chiariello, A.M.; Annunziatella, C.; Kraft, K.; Schöpflin, R.; Wittler, L.; Andrey, G.; Vingron, M.; Pombo, A.;, et al. Polymer Physics Predicts the Effects of Structural Variants on Chromatin Architecture. Nat. Genet. 2018, 50, 662–667, doi:10.1038/s41588-018-0098-8.

84. Chiariello, A.M.; Abraham, A.; Bianco, S.; Esposito, A.; Fontana, A.; Vercellone, F.; Conte, M.; Nicodemi, M. Multiscale Modelling of Chromatin 4D Organization in SARS-CoV-2 Infected Cells. Nat. Commun. 2024, 15, 4014, doi:10.1038/s41467-024-48370-6.

85. Fiorillo, L.; Musella, F.; Conte, M.; Kempfer, R.; Chiariello, A.M.; Bianco, S.; Kukalev, A.; Irastorza-Azcarate, I.; Esposito, A.; Abraham, A.;, et al. Comparison of the Hi-C, GAM and SPRITE Methods Using Polymer Models of Chromatin. Nat. Methods 2021, 18, 482–490, doi:10.1038/s41592-021-01135-1.

86. Fiorillo, L.; Bianco, S.; Chiariello, A.M.; Barbieri, M.; Esposito, A.; Annunziatella, C.; Conte, M.; Corrado, A.; Prisco, A.; Pombo, A.;, et al. Inference of Chromosome 3D Structures from GAM Data by a Physics Computational Approach. Methods 2020, 181–182, 70–79, doi:10.1016/j.ymeth.2019.09.018.

87. Conte, M.; Fiorillo, L.; Bianco, S.; Chiariello, A.M.; Esposito, A.; Nicodemi, M. Polymer Physics Indicates Chromatin Folding Variability across Single-Cells Results from State Degeneracy in Phase Separation. Nat. Commun. 2020, 11, 3289, doi:10.1038/s41467-020-17141-4.

88. Chiariello, A.M.; Bianco, S.; Oudelaar, A.M.; Esposito, A.; Annunziatella, C.; Fiorillo, L.; Conte, M.; Corrado, A.; Prisco, A.; Larke, M.S.C.;, et al. A Dynamic Folded Hairpin Conformation Is Associated with α-Globin Activation in Erythroid Cells. Cell Rep. 2020, 30, 2125–2135.e5, doi:10.1016/j.celrep.2020.01.044.

89. Conte, M.; Chiariello, A.M.; Bianco, S.; Esposito, A.; Abraham, A.; Nicodemi, M. Physics-Based Polymer Models to Probe Chromosome Structure in Single Molecules. In Methods in Molecular Biology; 2023; Vol. 2655.

90. Defenouillère, Q.; Yao, Y.; Mouaikel, J.; Namane, A.; Galopier, A.; Decourty, L.; Doyen, A.; Malabat, C.; Saveanu, C.; Jacquier, A.;, et al. Cdc48-Associated Complex Bound to 60S Particles Is Required for the Clearance of Aberrant Translation Products. Proc. Natl. Acad. Sci. U. S. A. 2013, 110, doi:10.1073/pnas.1221724110.

91. Joazeiro, C.A.P. Mechanisms and Functions of Ribosome-Associated Protein Quality Control. Nat. Rev. Mol. Cell Biol. 2019, 20.

92. Chu, J.; Hong, N.A.; Masuda, C.A.; Jenkins, B. V.; Nelms, K.A.; Goodnow, C.C.; Glynne, R.J.; Wu, H.; Masliah, E.; Joazeiro, C.A.P.;, et al. A Mouse Forward Genetics Screen Identifies LISTERIN as an E3 Ubiquitin Ligase Involved in Neurodegeneration. Proc. Natl. Acad. Sci. U. S. A. 2009, 106, doi:10.1073/pnas.0812819106.

93. Belaghzal, H.; Dekker, J.; Gibcus, J.H. Hi-C 2.0: An Optimized Hi-C Procedure for High-Resolution Genome-Wide Mapping of Chromosome Conformation. Methods 2017, 123, doi:10.1016/j.ymeth.2017.04.004.

94. Reiff, S.B.; Schroeder, A.J.; Kırlı, K.; Cosolo, A.; Bakker, C.; Lee, S.; Veit, A.D.; Balashov, A.K.; Vitzthum, C.; Ronchetti, W.;, et al. The 4D Nucleome Data Portal as a Resource for Searching and Visualizing Curated Nucleomics Data. Nat. Commun. 2022, 13, doi:10.1038/s41467-022-29697-4.

95. Dekker, J.; Belmont, A.S.; Guttman, M.; Leshyk, V.O.; Lis, J.T.; Lomvardas, S.; Mirny, L.A.; O’Shea, C.C.; Park, P.J.; Ren, B.;, et al. The 4D Nucleome Project. Nature 2017, 549, 219–226, doi:10.1038/nature23884.

96. De Gennes, P.G. Scaling Concepts in Polymer Physics. Cornell University Press. In Ithaca N.Y.,; 1979 ISBN 080141203X.

97. Shrinivas, K.; Sabari, B.R.; Coffey, E.L.; Klein, I.A.; Boija, A.; Zamudio, A. V.; Schuijers, J.; Hannett, N.M.; Sharp, P.A.; Young, R.A.;, et al. Enhancer Features That Drive Formation of Transcriptional Condensates. Mol. Cell 2019, 75, 549–561.e7, doi:10.1016/j.molcel.2019.07.009.

98. Esposito, A.; Bianco, S.; Chiariello, A.M.; Abraham, A.; Fiorillo, L.; Conte, M.; Campanile, R.; Nicodemi, M. Polymer Physics Reveals a Combinatorial Code Linking 3D Chromatin Architecture to 1D Chromatin States. Cell Rep. 2022, 38, 110601, doi:10.1016/j.celrep.2022.110601.

99. Thompson, A.P.; Aktulga, H.M.; Berger, R.; Bolintineanu, D.S.; Brown, W.M.; Crozier, P.S.; in ’t Veld, P.J.; Kohlmeyer, A.; Moore, S.G.; Nguyen, T.D.;, et al. LAMMPS - a Flexible Simulation Tool for Particle-Based Materials Modeling at the Atomic, Meso, and Continuum Scales. Comput. Phys. Commun. 2022, 271, 108171, doi:10.1016/j.cpc.2021.108171.

100. Plimpton, S. Fast Parallel Algorithms for Short-Range Molecular Dynamics. J. Comput. Phys. 1995, 117, 1–19, doi:10.1006/jcph.1995.1039.

101. Conte, M.; Esposito, A.; Fiorillo, L.; Campanile, R.; Annunziatella, C.; Corrado, A.; Chiariello, M.G.; Bianco, S.; Chiariello, A.M. Efficient Computational Implementation of Polymer Physics Models to Explore Chromatin Structure. Int. J. Parallel, Emergent Distrib. Syst. 2019, doi:10.1080/17445760.2019.1643020.

102. Kremer, K.; Grest, G.S. Dynamics of Entangled Linear Polymer Melts: A Molecular-Dynamics Simulation. J. Chem. Phys. 1990, 92, 5057–5086, doi:10.1063/1.458541.

103. Allen, M.P.; Tildesley, D.J. Computer Simulation of Liquids (Oxford Science Publications) SE - Oxford Science Publications. Oxford Univ. Press 1989.

104. Conte, M.; Fiorillo, L.; Annunziatella, C.; Esposito, A.; Musella, F.; Abraham, A.; Bianco, S.; Chiariello, A.M. Dynamic and Equilibrium Properties of Finite-Size Polymer Models of Chromosome Folding. *Phys*. Rev. E 2021, 104, 054402.

105. Baum, M.; Erdel, F.; Wachsmuth, M.; Rippe, K. Retrieving the Intracellular Topology from Multi-Scale Protein Mobility Mapping in Living Cells. Nat. Commun. 2014, 5, 4494, doi:10.1038/ncomms5494.

106. Conte, M.; Chiariello, A.M.; Abraham, A.; Bianco, S.; Esposito, A.; Nicodemi, M.; Matteuzzi, T.; Vercellone, F. Polymer Models of Chromatin Imaging Data in Single Cells. Algorithms 2022, 15, 330, doi:10.3390/a15090330.

107. Bishop, M.; Michels, J.P.J. Polymer Shapes in Three Dimensions. J. Chem. Phys. 1986, 85, doi:10.1063/1.451508.

108. Coniglio, A.; Nicodemi, M. A Statistical Mechanics Approach to the Inherent States of Granular Media. Phys. A Stat. Mech. its Appl. 2001, 296, doi:10.1016/S0378-4371(01)00190-X.

109. Nicodemi, M. Force Correlations and Arch Formation in Granular Assemblies. Phys. Rev. Lett. 1998, 80, 1340–1343, doi:10.1103/PhysRevLett.80.1340.

110. Nicodemi, M.; Coniglio, A. Macroscopic Glassy Relaxations and Microscopic Motions in a Frustrated Lattice Gas. *Phys*. Rev. E 1998, 57, R39–R42, doi:10.1103/PhysRevE.57.R39.

111. Piegari, E.; Cataudella, V.; Di Maio, R.; Milano, L.; Nicodemi, M. A Cellular Automaton for the Factor of Safety Field in Landslides Modeling. Geophys. Res. Lett. 2006, 33, doi:10.1029/2005GL024759.

112. Oliveira, L.P.; Jensen, H.J.; Nicodemi, M.; Sibani, P. Record Dynamics and the Observed Temperature Plateau in the Magnetic Creep-Rate of Type-II Superconductors. Phys. Rev. B - Condens. Matter Mater. Phys. 2005, 71, doi:10.1103/PhysRevB.71.104526.

113. Nicodemi, M.; Fierro, A.; Coniglio, A. Segregation in Hard-Sphere Mixtures under Gravity. An Extension of Edwards Approach with Two Thermodynamical Parameters. Europhys. Lett. 2002, 60, 684–690, doi:10.1209/epl/i2002-00363-0.

114. Hamon, D.; Nicodemi, M.; Jensen, H.J. Continuously Driven OFC: A Simple Model of Solar Flare Statistics. Astron. Astrophys. 2002, 387, doi:10.1051/0004-6361:20020346.

115. Arenzon, J.J.; Nicodemi, M.; Sellitto, M. Equilibrium Properties of the Ising Frustrated Lattice Gas. J. Phys. I 1996, 6, doi:10.1051/jp1:1996120.

116. Cataudella, V.; Franzese, G.; Nicodemi, M.; Scala, A.; Coniglio, A. Critical Clusters and Efficient Dynamics for Frustrated Spin Models. Phys. Rev. Lett. 1994, 72, doi:10.1103/PhysRevLett.72.1541.

117. Borrelli, A.; De Falco, I.; Della Cioppa, A.; Nicodemi, M.; Trautteur, G. Performance of Genetic Programming to Extract the Trend in Noisy Data Series. Phys. A Stat. Mech. its Appl. 2006, 370, doi:10.1016/j.physa.2006.04.025.

118. Caglioti, E.; Coniglio, A.; Herrmann, H.J.; Loreto, V.; Nicodemi, M. Segregation of Granular Mixtures in the Presence of Compaction. Europhys. Lett. 1998, 43, doi:10.1209/epl/i1998-00402-x.

119. Tarzia, M.; Candia, A. de; Fierro, A.; Nicodemi, M.; Coniglio, A. Glass Transition in Granular Media. Europhys. Lett. 2004, 66, 531–537, doi:10.1209/epl/i2004-10015-y.

120. Iannone, F.; Ambrosino, F.; Bracco, G.; De Rosa, M.; Funel, A.; Guarnieri, G.; Migliori, S.; Palombi, F.; Ponti, G.; Santomauro, G.;, et al. CRESCO ENEA HPC Clusters: A Working Example of a Multifabric GPFS Spectrum Scale Layout. In Proceedings of the 2019 International Conference on High Performance Computing & Simulation (HPCS); IEEE, July 2019; pp. 1051–1052.

